# Functional grading of pericellular matrix surrounding chondrocytes: potential roles in signaling and fluid transport

**DOI:** 10.1101/365569

**Authors:** F. Saadat, M.J. Lagieski, V. Birman, S. Thomopoulos, G.M. Genin

## Abstract

The extracellular matrix surrounding chondrocytes within cartilage and fibrocartilage has spatial gradients in mechanical properties. Although the function of these gradients is unknown, the potential exists for cells to tailor their mechanical microenvironment through these gradients. We hypothesized that these gradients enhance fluid transport around the cell during the slow loading cycles that occur over the course of a day, and that this enhancement changes the nature of the mechanical signals received at the surface of the cell. To test this hypothesis, we studied the effect of these gradients on the mechanical environment around a chondrocyte using a closed form, linearized model. Results demonstrated that functional grading of the character observed around chondrocytes in articular cartilage enhances fluid transport, and furthermore inverts compressive radial strains to provide tensile signals at the cell surface. The results point to several potentially important roles for functional grading of the pericellular matrix.

## 1 Introduction

Chondrocytes within cartilage and fibrocartilage live in a poorly vascularized, nutrient-deprived extracellular matrix (ECM)(1; 2). Their mechanical microenvironment varies both with their location within the tissue and with the duration and time history of mechanical loading. The macroscopic mechanics of cartilage has long been known to be dominated by the poroelastic interaction of charged fluid flow with electrostatic and hydraulic resistance of the polyanionic proteoglycans within the ECM (3; 4). This behavior is well modeled by the standard biphasic model of cartilage that maps how fluid flow determines the time variation of cartilage mechanical response between the extremes of very fast loading (under a second, when fluid is pressurized and carries most of the stress) and very slow loading (hours, when fluid flow has ceased and stress is carried by the ECM) (5; 6). In both of these extremes, the largely elastic ECM response varies from uniaxial compression in the superficial zone (where the surface is free to deform under traction) to hydrostatic compression in the deeper zones (where the tissue is prevented from deforming by the rigid bone).

At the level of individual chondrocytes, these fields are modulated by the nature of the pericellular ECM (PCM) surrounding the cell. The PCM differs from ECM far from the cell in its composition (7; 8), structure (9; 10), and mechanical properties (11–17). Mechanical effects of this PCM are readily observed: although chondrocytes are compliant (elastic modulus 0.1 – 8 kPa, (18–24)) compared to the surrounding ECM (elastic modulus 0.1 – 2 MPa (17; 25; 26)), they can deform *less* than the ECM when cartilage is compressed (27–30). This suggests that the PCM, which envelops the chondrocyte to form a chondron, shields the cell from large deformation (31–34). Numerical models (35–38) and microscopy studies (30; 39) of this shielding reveal that the PCM is important to cell function, and that its contributions depend upon the relative mechanical properties of the cell, PCM and ECM. Reported elastic moduli of the PCM vary over two orders of magnitude, from 1.5 to 162 kPa, but are typically intermediate to moduli of the chondrocytes and ECM (40; 41).

More recent AFM mapping reveals a monotonic grading of modulus from that of the peri-cellular region just outside the cell membrane to that of the ECM (15; 16; 42). Our objective was to characterize how this functional grading of mechanical properties affects the cell microenvironment and the mechanical shielding of chondrocytes. Our focus was the conditions of slow or fast loading, where elasticity dominates, and also the deep zone of cartilage, where the stress field approaches hydrostatic compression.

Two challenges surround the function of chondrocytes in the deep zone of cartilage. The first is receiving mechanical cues. Mechanical shielding by the PCM is likely important for blocking the pathological effects of stretch-activated channels such Piezo1, Piezo2, and TRPV4 (43; 44), and for avoiding the associated degenerative pathologies such as osteoarthritis (OA) (45). However, mechanical shielding might be undesirable in the deep zone given the important roles of mechanical loading in cartilage health and chondrocyte metabolism (2; 35; 46–48), and given the presence of physiological protein structures such as integrin plaques that require tensile strains (49; 50). We therefore asked whether functional grading of the PCM might be tailored to tune the tensile signals that chondrocytes in the deep zone receive. The second challenge is nutrient transport. Given the relatively attenuated fluid flow available to deliver nutrients to chondrocytes in the deep zone, is it possible that functional grading of the PCM might be tailored to amplify fluid exchange in the PCM?

Our approach was to study simplified theoretical models appropriate to slow and fast loading, and appropriate to the hydrostatic conditions of the deep zone. We explored how the modulus mismatch and functional grading between the chondrocyte and the ECM can enhance fluid transport and can enable the phenomenon of strain inversion, in which a compressive loading cycle can yield tensile radial strains at the periphery of a chondrocyte. Results suggested that functionally graded PCM, as reported by Wilusz et al. (15) for both healthy and OA cartilage, can enhance pericellular dilatation and induce strain inversion at the cell periphery during a compressive loading.

## 2 Methods

We first established the degree to which uniaxial compressive stress is expected to lead to hydrostatic compressive stress in the deep zone of articular cartilage. Because the stress field in cartilage shifted from uniaxial in the superficial layer to nearly hydrostatic in the deep layer, we studied the effect of functionally graded PCM in response to hydrostatic loading.

The mechanical microenvironment of a chondrocyte was assessed through a simple mathematical model of a chondrocyte surrounded by PCM and embedded within an infinite ECM. The system of a cell, its PCM, and the ECM was idealized as three concentric spherical layers including (Figure 1): (1) a central chondrocyte of radius of *r*_0_, (2) a graded PCM with an outer radius *r*_*N*_, and (3) an outer sheath of ECM of infinite radius. This initial spherical shape of the chondrocyte and PCM is consistent with microscopic observation (28; 42; 51). For the purpose of our qualitative analysis, linear material properties and small strain conditions were studied. Although cells, PCM and ECM are in reality both nonlinear and viscoelastic, the linear elastic idealization studied here serves to identify behaviors that can be expected under monotonic loading at a rate that is either rapid or slow compared to viscoelastic relaxation spectra of the ECM or PCM.

**Figure 1:**
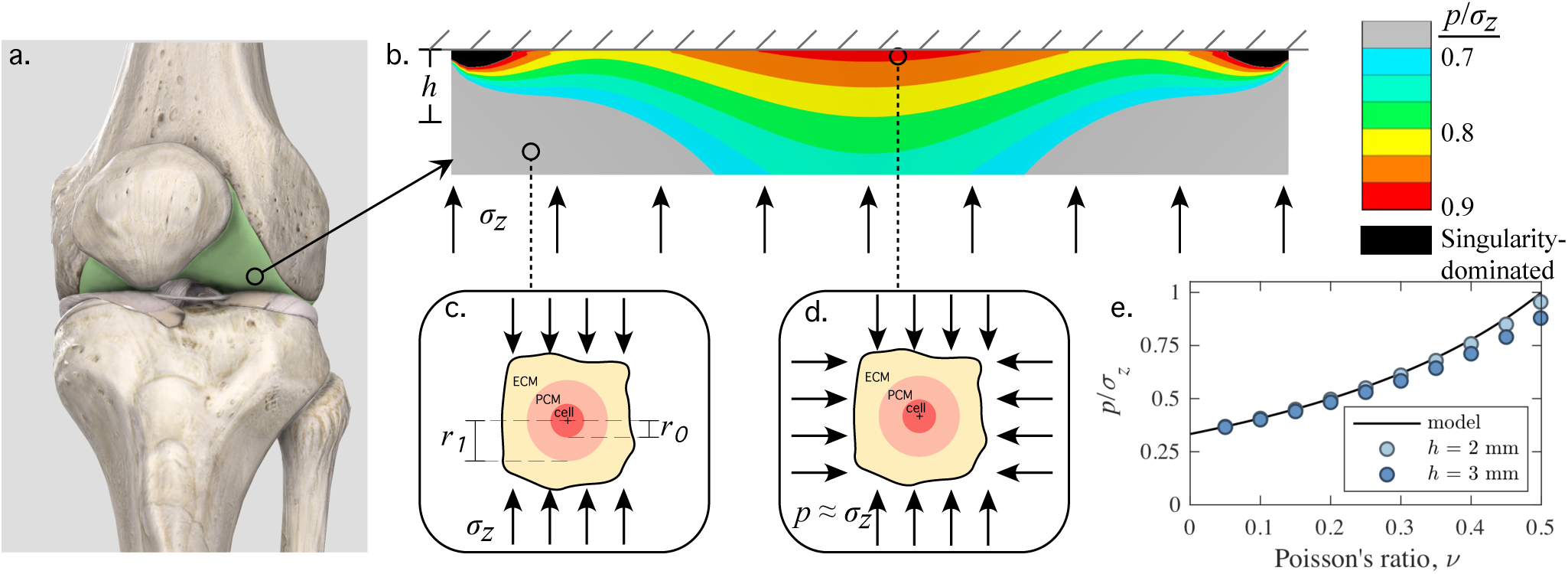
Cartilage (a) in the deep zone can be expected to experience high levels of compressive hydrostatic loading. (b) Illustration of the contours of hydrostatic stress for Poisson’s ratio *v* = 0.49. (c) In the superficial zone, the stress field is well approximated by uniaxial compression. (d) In the deep zone, the stress field approached hydrostatic compression. (e) The model and simulation show that the degree of hydrostatic compression is a function of Poisson’s ratio. Panel (a) was generated using the Complete Anatomy software package (3D4Medical, Ltd., Dublin, Ireland.)

To quantify the mechanical response of this system to a radial compressive stress *p* (Figure 1d), the system was analyzed using an exact theory of the elasticity solution, described in detail in the Appendices. The cell, PCM, and ECM were each treated as isotropic with two independent material constants: a Young’s modulus (*E*_*ECM*_) and a Poisson ratio (*v*_*ECM*_) for the ECM; corresponding values *E*_*cell*_ and *v*_*cell*_ for the cell; and a gradation in the PCM that was modeled as an assembly of *N* concentric and perfectly bonded piece-wise constant isotropic spherical shells. In the following, we describe the analysis, including two associated boundary value problems that were studied to better interpret the mechanical environment of the chondrocyte and its behavior during compression.

### 2.1 Transition from uniaxial to hydrostatic compression in a cartilage layer

At the central region just below the articular surface of compressed cartilage, where shear stresses associated with flow of synovial fluid are small, the surface traction is approximated reasonably by uniaxial compression (Figure 1b). To gain a sense of the degree to which this uniaxial traction converts to hydrostatic compression near the osteochondral interface, a finite element analysis was performed, and a closed form approximation was developed. In both cases, the focus was loading over time intervals that were either long or short compared to the characteristic relaxation times, so that separation of time scales ensured that linear elasticity served as a reasonable first approximation.

For the finite element analyses, a disc of radius 10 mm and thickness of either 2 mm or 3 mm was compressed with a vertical traction on one circular face, and held fixed on the other. Because of the contrast with the rigid constraint, the Youngs modulus does not affect the stress distribution in such a setting, but the Poisson ratio affected the degree to which hydrostatic stresses arose at the interface, resulting in the emergence of a Williams-type singularity at the free edge of the cartilage-bone interface (52; 53).The Poisson ratio was thus varied, and the hydrostatic stress averaged over a region of 100 micrometers at the center of the disc was calculated and compared to the analytical estimate. For all simulations, quadratic interpolation quadrilateral elements were used. Standard convergence studies showed that analyses using the final mesh, with 48,962 degrees of freedom, had converged to within a few percent outside of the Williams-type free edge singularity. Equations were solved using the ANSYS software system (Ansys, Canonsburg, PA).

To estimate the peak hydrostatic stress in closed form, a simple boundary value problem was studied. At the loaded surface, the stress was uniaxial, The compressive strain in the direction of compression was 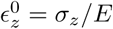 at the surface, and those in the radial and tangential directions were 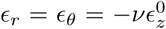, where *v* and *E* Poisson ratio and elastic modulus, respectively, for either fast or slow loadings.

The tissue in the deep zone is stiffened against compression by the lateral constraint of bone. This effect is well known as the classic Boussinesq problem of linear elasticity, it is also present in the classic frictional punch problem (54). Assuming perfect adhesion between the cartilage and bone, neglecting functional grading for the purpose of this illustration, and invoking Saint Venants principle (54) to disregard the Williams-type free edge singularity, a reasonable closed-form approximation to the stress and strain can be written by setting *ϵ*_*r*_ = *ϵ*_*θ*_:

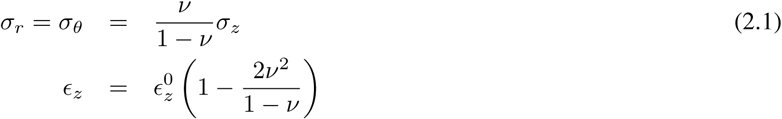

For an isotropic material, *ϵ*_*z*_ at the base is attenuated by the term 2*v*^2^/(1 – *v*), and the stress state at that point approaches hydrostatic compression depending upon Poisson’s ratio. The hydrostatic pressure *p* is one third the trace of the stress tensor:

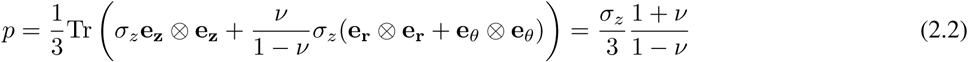

For *v* = 1/2, the compressive strain is fully attenuated, and the stress state is fully hydrostatic.

### 2.2 Models of a cell embedded in a compressed ECM

We began by studying a chondrocyte in the absence of PCM, both because this problem was of interest as an idealized comparison case and because it might be relevant at certain stages of development (55–57). We conducted a theoretical analysis on a two-layer model including an isolated cell in a uniform ECM and investigated how a broad range of possible values for the mechanical properties of cell and ECM influenced the mechanical environment of the individual cell. The stress and displacement fields for this case are derived in Appendices A, B, and C.

In the second step of our analysis, we included a PCM layer in one of two ways. The first was through a functionally-graded PCM based upon the experimental data of Wilusz et al. (15). Here, the Poisson ratio of the PCM was held constant (*v*_*PCM*_ = 0.04), while the elastic modulus was graded. Wilusz et al. (15) report an abrupt change of modulus at cell-PCM interfaces, and this was captured in our simulations.

Note that a wide range of Poisson ratios has been reported for the ECM and PCM (17), and that the abrupt jump in elastic modulus might be due to difficulties in measuring the stiffness very close to the cell-PCM interface. We therefore evaluated this by studying simplified models. Here, both the elastic modulus (*E*_*PCM*_), and Poisson ratio (*v*_*PCM*_) were varied smoothly from those of the cell to those of the ECM. The full range of monotonic functional gradings was studied by modulating the exponent, *k*, of a power law interpolation function (Figure 6): for *k* = 0, the PCM had the same mechanical properties as the ECM; for *k* = ∞, the PCM had the mechanical properties of the cell; for *k* = 1, the interpolation was linear.

All equations were solved using custom written codes in Mathematica (Wolfram Research, Champaign, IL) and Matlab (The Mathworks, Natick, MA).

## 3 Results and Discussion

### 3.1 The stress field in cartilage varied with depth from uniaxial compression to nearly hydrostatic compression

At small strain and over a time interval that is either long or short compared to its characteristic relaxation times, the stress field in the superficial zone was well approximated by uniaxial compression (Figure 1). However, the tissue in the deep zone was stiffened against compression by the lateral constraint of bone, and the stress field shifted to hydrostatic compression in a way that was modulated by Poisson’s ratio and that was well-predicted by the model in equation 2.2. The hydrostatic constraint died out towards the periphery of the elastic model; Williams-type free edge singularities were evident at the constrained corners. However, in the deep zone the stress field was close to hydrostatic.

### 3.2 Tensile strains can arise at the cell periphery during hydrostatic compression

We began analysis of the system by considering the comparison case of a two-layer model of an isolated cell within an infinite uniform ECM. When this model was loaded in hydrostatic compression, the radial strain field at the periphery of the ECM inverted: although the radial strain infinitely far from the cell was compressive, the radial strain at the cell periphery was tensile (Figures 2 and 3). Although strain inversion has been reported earlier in a finite-element model for a chondrocyte within an ECM (35), it has never been explored in detail, and we therefore explored how the amplitude and spatial extent of this strain inversion depended upon material properties. For a cell with Young’s modulus *E*_*cell*_ = 0.65 kPa and Poisson’s ratio *v*_*cell*_ = 0.38 (22; 58) embedded within an ECM with *E*_*ECM*_ = 306 MPa (17), the amplitude and spatial extent of the strain inversion varied with the Poisson ratio of the ECM (Figure 2). The values of *v*_*ECM*_ = 0.04 and *v*_*ECM*_ = 0.38 chosen for illustration are within the range reported in the literature (59–62).

**Figure 2:**
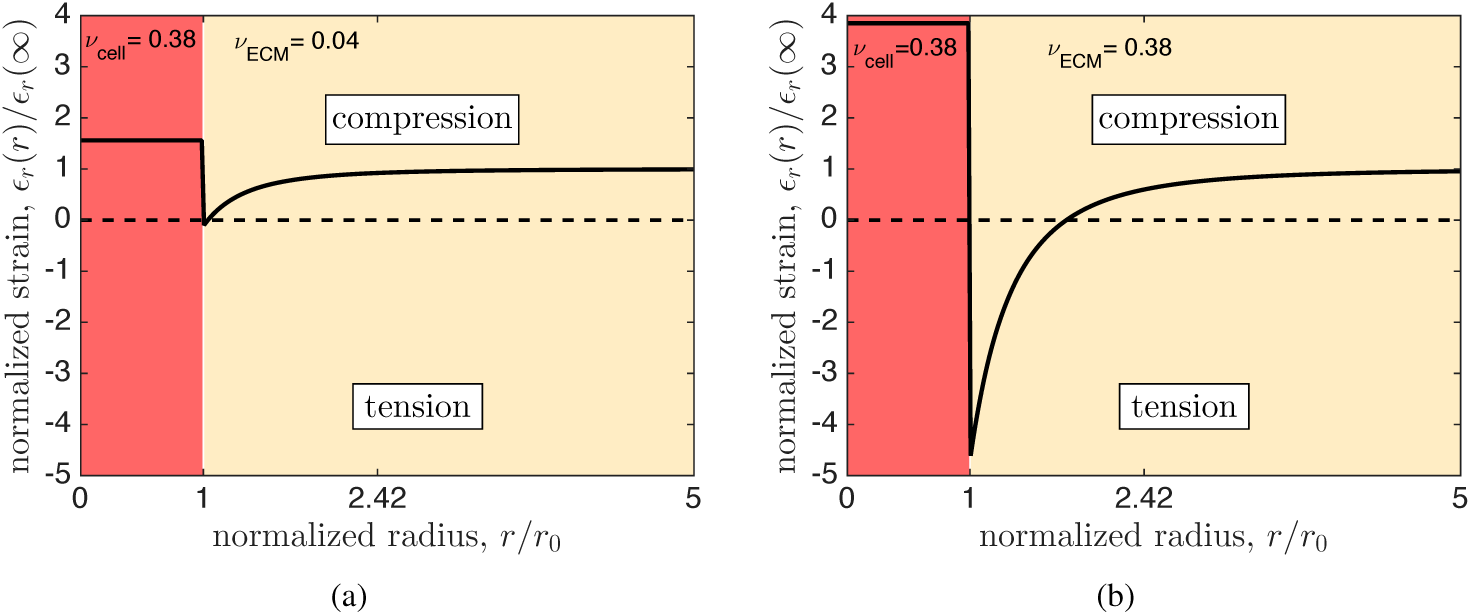
The radial strain at the periphery of a cell is tensile in response to hydrostatic loading of the ECM in a simplified model of a cell embedded directly within the ECM. The amplitude and spatial extent of the strain inversion increased with increasing Poisson ratio *v*_*ECM*_ of the extracellular matrix. (a) *v*_*ECM*_ =0.04; (b) *v*_*ECM*_ =0.38.

### 3.3 A phase diagram suggests that strain inversion occurs in human cartilage

To explore how the full range of possible moduli and Poisson’s ratios would affect strain inversion, a full parametric study was performed, and a phase diagram was generated. Both the amplitude (Figure 3a) and size (Figure 3b) of the inversion zone showed a nonlinear phase boundary. Strain inversion was impossible for properties that lay below the solid black phase boundary (i.e., for properties in the yellow region). Strain inversion always occurred for sufficiently high Poisson ratio and sufficiently stiff ECM. When evaluating material properties associated with both healthy and osteoarthritic human cartilage (gray and white circles, Figure 3), the properties for these cases lay well within the portion of the phase diagram in which strain inversion occurs. The degree of strain inversion was robust against changes associated with osteoarthritis. However, we note that the effects of osteoarthritis on Poisson contraction of a chondrocyte have yet to be measured.

**Figure 3:**
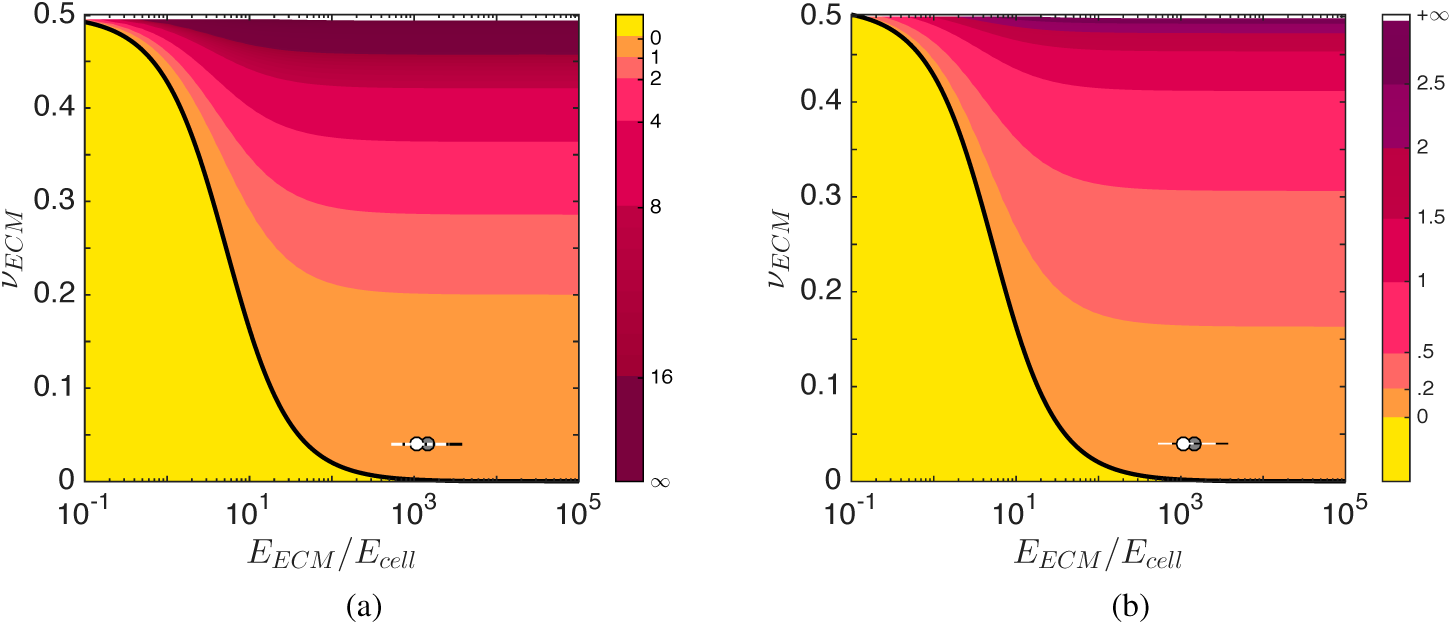
Phase diagrams for strain inversion at the periphery of a cell embedded in a uniform ECM. (a) Amplitude of the strain inversion, *ϵ*̄ = *ϵ*_*r*_ (*r* = *r*_0_)/*ϵ*_*r*_ (∞). (b) Thickness of the strain inversion zone (*l*/*r*_0_). The gray and white circles represent properties for normal and osteoarthritic conditions in human articular cartilage. The degree of strain inversion was robust against changes associated with osteoarthritis. For these plots, Poisson’s ratio of the cell was held at *v*_*cell*_ =0.38.

The properties of cells and ECM used in Figure 3 represent relatively slow loading of cartilage (15; 22; 36). For loading sufficiently fast that water flow is not significant, the Poisson ratio of the ECM increases, and the amplitude and spatial extent of strain inversion increase as well. These findings have implications for study of chondrocytes *in vitro* (63–66) as they suggest that the stiffening the ECM has the surprising consequence of amplifying tensile radial strain at the cell periphery.

### 3.4 The PCM enhances fluid transport at the cell periphery, while preserving strain inversion

The next simplified problem studied before evaluating a realistic PCM was a three-layered model of an elastic cell surrounded by a homogenous PCM layer that was in turn surrounded by a homogenous ECM layer (Figure 4). The radius of the PCM region was set to 2.42 times the cell radius, following the observations of Alexopoulos et al. (12). For this illustration, we again chose we again chose *E*_*cell*_ = 0.65*kPa* and *E*_*ECM*_ = 306 kPa (17), *v*_*cell*_ = 0.38 (58), and *v*_*ECM*_ = 0.04 (13). Following Kim et al. (13), we chose *E*_*PCM*_ = 104 kPa. Following Darling et al. (17), we assigned *v*_*ECM*_ = *v*_*PCM*_.

**Figure 4:**
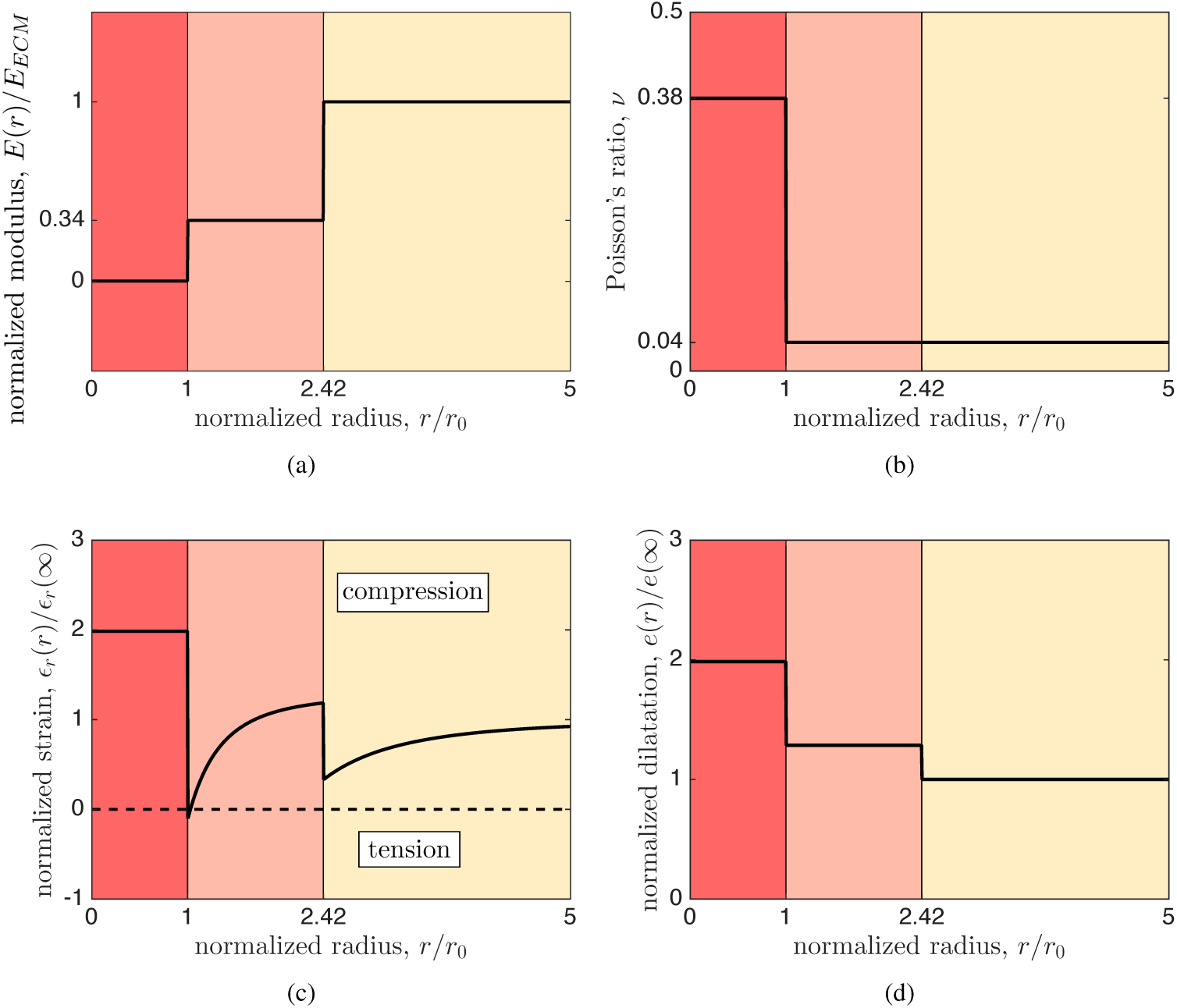
Strain inversion and enhanced volumetric cycling in the PCM around a chondrocyte. (a and b) As a first comparison study, a uniform elastic modulus in the PCM was studied, with magnitude as reported by (17). Dimensions were as reported by Jones et al. (22). (c) For this model problem, the normalized radial strain was inverted at the cell periphery. (d) The PCM served to enhance the volumetric change in a loading cycle: the cell and PCM experienced enhanced dilatation compared to the remote ECM.

For this case, a relatively small strain inversion zone was observed at the cell periphery, which would serve to insulate the cell membrane from compressive radial strain and to provide mild tensile stimulus during compression (Figure 4c). This layer served an additional function in that it enhanced the volumetric change in a loading cycle: the cell and PCM experienced enhanced dilatation compared to the remote ECM (Figure 4d). This suggests that, over the course of a slow daily loading cycle of cartilage, in which water flow is dominant, the mechanics of a PCM region can be tuned to amplify the fluid transport into and out of the pericellular region.

To examine the degree to which this enhanced dilatation and strain inversion are present in healthy and osteoarthritic human cartilage, a functionally graded PCM was studied with the elastic modulus following the spatial grading reported by Wilusz et al. (15) (Figure 5a). In osteoarthritic cartilage, this grading began with a slightly elevated stiffness at the cell periphery, and then rose gradually until the edge of the PCM to the value in the ECM. Note that only *relative* moduli factor into the solution. Although healthy and osteoarthritic tissues have substantially different moduli, their gradients in modulus, when normalized to their ECM moduli, are nearly identical (Figure 5a). Results indicate strain inversion and amplified volumetric expansion of a magnitude similar to that observed in the idealized test case having constant modulus within the PCM (Figures 5c and 5d), and were similar for both healthy and osteoarthritic (OA) tissues.

**Figure 5:**
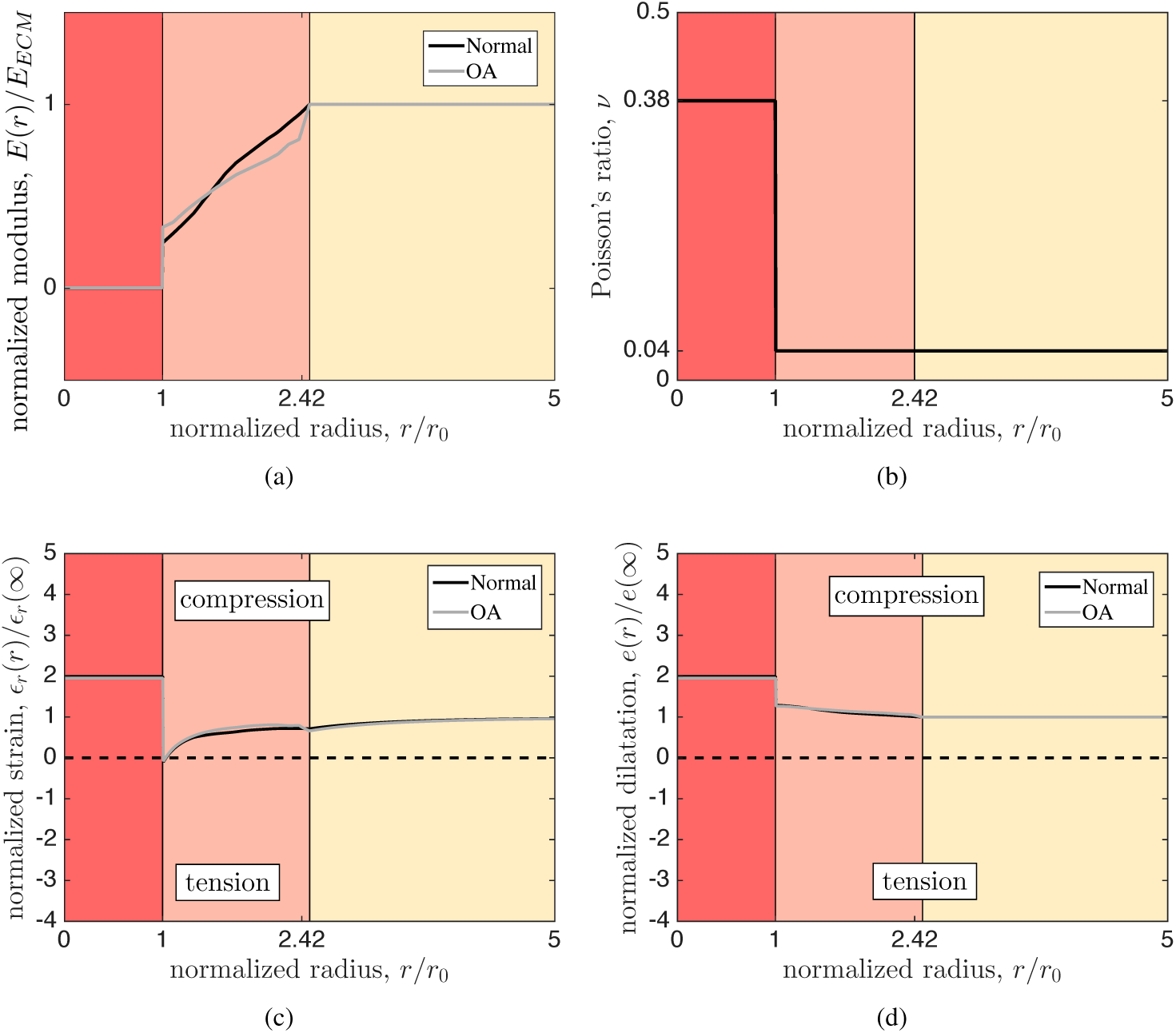
Strain inversion zone and enhanced dilatation around an idealized chondrocyte. (a) Here, the grading of elastic moduli reported by (15) was applied in conjunction with (b) constant Poisson ratios within each layer (*v*_*cell*_ = 0.38 (58), *v*_*PCM*_ = *v*_*ECM*_ = 0.04 (13)). (c) A relatively small strain inversion region was predicted, as well as (d) a graded region of enhanced PCM dilatation, of magnitude slightly greater than that predicted in the absence of grading (cf. Fig. 2). The effects appear robust against changes associated with OA. The dimensions reported by (15), used to generate this figure, differ slightly from those used in all other figures in this document: *r*_0_ = 5 *μ*m and *r*_*N*_ = 12.5 *μ*m.

**Figure 6:**
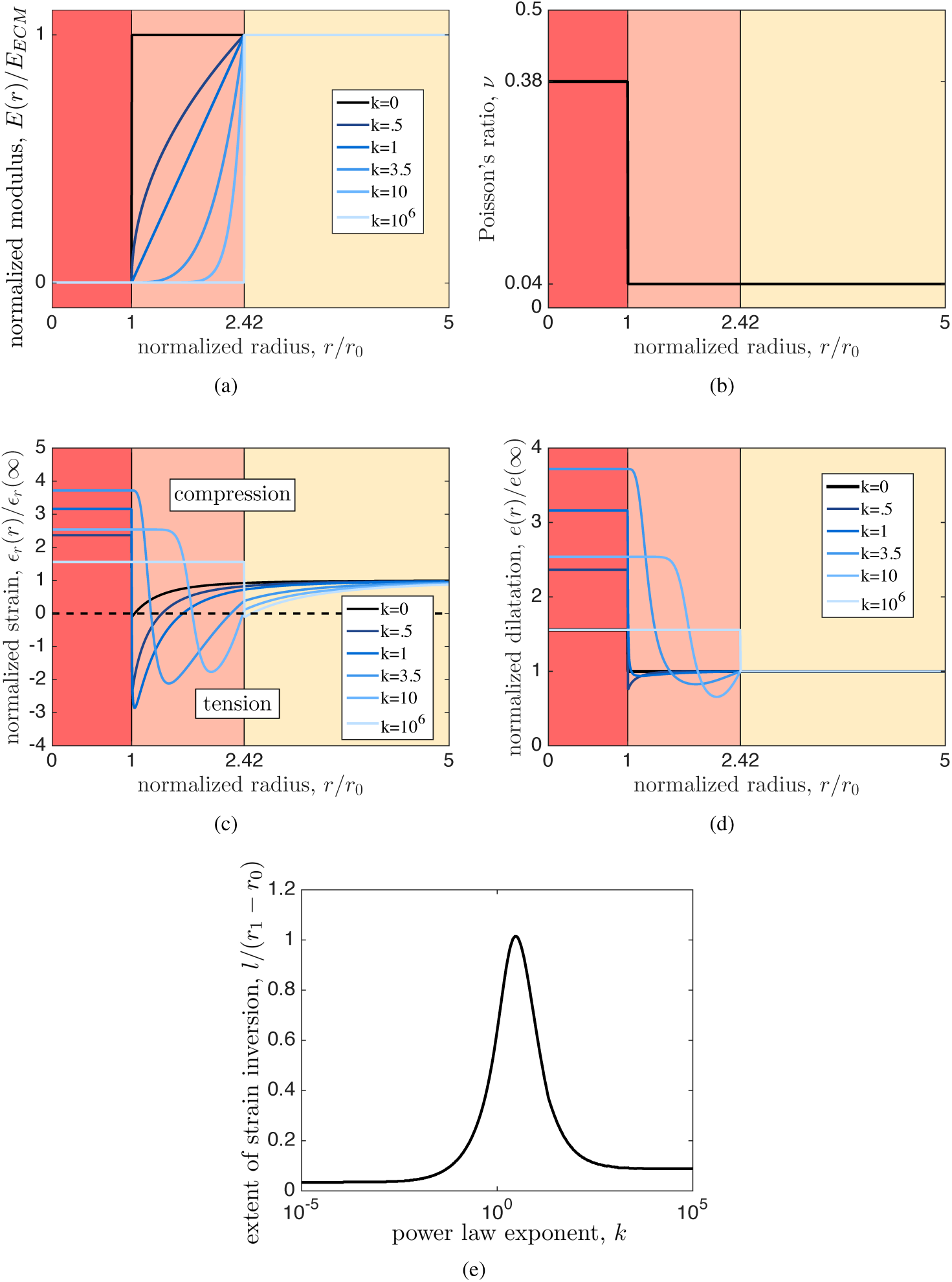
The size, extent, and character of the strain inversion zone can be tailored though the nature of the interpolation of mechanical properties though the PCM. (a and b) Modulus grading was studied by varying the power law exponent *k* of the interpolation function, while the Poisson ratio of the PCM was set to that of the ECM. (c) Functional grading of the PCM controlled the extent and positioning of the strain inversion zone with respect to the cell boundary, and could increase the magnitude of the inverted radial strain at the chondrocyte-matrix interface to nearly four times the nominal strain in the remote ECM. (d) Dilatation of the cell and PCM could be increased similarly by functional grading. (e) Parametric analysis revealed that the relative size of the strain inversion zone was maximized for a value of *k* associated with interpolation that was slightly concave up.

### 3.5 Functional grading can be harnessed to tailor strain inversion and dilatation enhancement

To evaluate the degree to which strain inversion and dilatation can be tailored by functional grading of PCM modulus, a parametric study was performed over a broad range of smooth, monotonic gradings (Figure 6). For this, the functional grading of the PCM was set to follow a power law of exponent *k*:

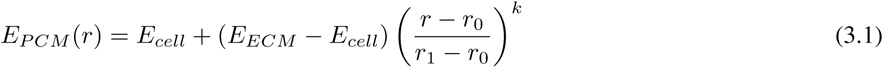

where *r*_*0*_ is the radius of the cell and *r*_1_ is the outer radius of the PCM. For *k* = 0, the PCM matched the ECM; for *k* → ∞, the PCM matched the cell; for *k* = 1, the interpolation was linear (Figures 6a and 6b). Functional grading of the PCM controlled the extent and positioning of the strain inversion zone with respect to the cell boundary, and could increase the magnitude of the inverted radial strain at the chondrocyte-matrix interface to nearly four times the nominal strain in the remote ECM (Figure 6c). Dilatation of the cell and PCM could be increased similarly by functional grading (Figure 6d). These findings show that functional grading of the PCM can eliminate the stress concentration that might otherwise arise at the cell interface due to the material mismatch (53; 67). Parametric analysis, described in Appendix B, revealed that the relative size *l* of the strain inversion zone was maximized for a value of *k* associated with interpolation that was slightly concave up (Figure 6e). At this maximum, strain inversion occurred across the entire PCM.

### 3.6 Caveats

We note that real cells and ECM are nonlinear and viscoelastic or poroelastic (23; 42; 68–70), and that even for the linear models, specifying the appropriate material parameters is a largely unmet challenge. However, these results show the potential for a chondrocyte in the deep zone of articular cartilage to tailor its mechanical microenvironment through functional grading of the PCM. Results motivate additional imaging analysis of compressed cartilage to identify whether fluid flow around the cell is indeed enhanced. Results show that, at least for small loading increments, PCM grading can be expected to affect mechanical signaling at the cell membrane.

## 4 Conclusion

The models studied in this work suggest that functional grading of PCM mechanics can enable a chondrocyte to tailor the mechanical stimuli it receives and to tailor volumetric flow at the cell periphery. Although the degree of confidence in published material properties, especially Poisson’s ratios, were not sufficient to identify the degree to which this might occur *in vivo*, the values that are available suggested that this is at least a possible scenario. Results motivate further study of the detailed mechanical properties ad strain fields around chondrocytes, and suggest a new pathway by which remodeling of the cell microenvironment can tailor cellular mechanotransduction.

## 5 Author contributions

All authors participated in design of the research; F.S. and M.L. performed the research; all authors contributed to the writing of the manuscript.

## 6 Acknowledgments

The authors thank Farshid Guilak for critical comments on the manuscript. This work was funded in part by the National Institutes of Health through grant U01EB016422, and by the National Science Foundation through the Science and Technology Center for Engineering Mechanobiology, grant CMMI 1548571.

## A Mechanics of a spherical cell within a functionally graded PCM

We approximated a functionally graded solid by considering a system of concentric elastic, isotropic spherical layers surrounding an elastic, isotropic spherical cell. When loaded by a radial compressive stress *p* applied at the outer boundary, the stress field in layer *i*, in spherical (*r*, *θ*, *ϕ*) coordinates is (e.g. (71)):

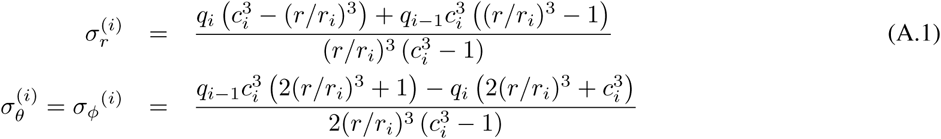

where *q*_*i*_ and *q*_*i* – 1_ are the the radial compressive stresses at the outer and inner interface, respectively, *r* is the radial coordinate, *c*_*i*_ = *r*_*i* – 1_/*r*_*i*_ is the ratio of the inner radius *r*_*i* – 1_ to the outer radius *r*_*i*_, and all shear stresses are zero. The displacement field is:

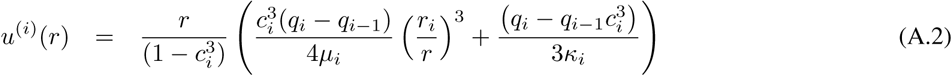

where *κ*^(*i*)^ is the bulk modulus and *ν*^(*i*)^ is Poisson’s ratio of layer *i*. The stresses *q*_*i*_ were found by considering continuity of displacement at the interfaces between adjacent layers, *u*^(*i*)^(*r* = *r*_*i*–1_) = *u*^(*i*–1)^(*r* = *r*_*i*–1_), *i* = 1,2,…*N*. This yielded the recurrence relation:

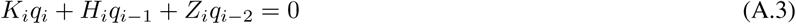

where the coefficients are

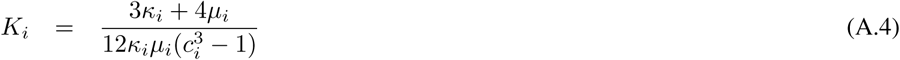

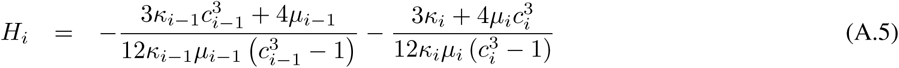

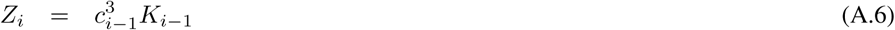

For *N* concentric layers encompassing the solid elastic spherical cell (*i*=0), the number of continuity conditions in Equation (A.3) is *N* – 1, but the number of unknown *q*_*i*_ is *N* + 1. The remaining two equations are equilibrium along the outermost and spherical layer (radius *r*_*N*_) and innermost sphere (radius *r*_0_). For the former, *q*_*N*_ = *p*. For the latter, we note that the stress field inside the cell is hydrostatic and uniform, with 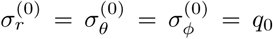. Accordingly, the strain field is 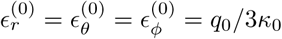, and the displacement at the boundary is:

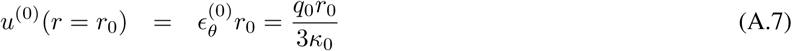

where *κ*_0_ is the bulk modulus of the cell. The continuity requirement along the outer edge of the cell is thus:

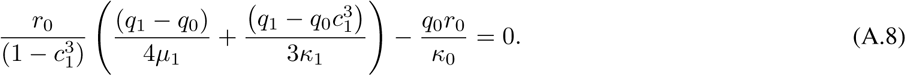

All interfacial radial stresses can then be determined from Equations A.3 and A.8, and the relation *q*_*N*_ = *p*.

## B Scaling of the strain inversion zone

To assess how the size and amplitude of the strain inversion zone scale with mechanical parameters, we studied a spherical cell of radius *r*_0_ in an infinite ECM (radius *r*_*N*_ → ∞) with no PCM. The magnitude of strain inversion scales as:

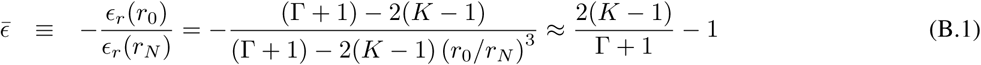

where *K* = *κ*_*ECM*_/*κ*_*cell*_ and Γ = (4/3)(*μ*_*ECM*_/*κ*_*cell*_), in which *κ*_*cell*_ and *κ*_*ECM*_ represent the bulk moduli of the cell and ECM, respectively, and *μ*_*ECM*_ is the shear modulus of the ECM. The approximation represents the limit as *r*_*N*_ → ∞.

The thickness of the strain inversion zone was calculated by finding the roots of the strain function that obtained from the equation representing the strain as a function of the position. The thickness of the strain inversion zone, normalized by the cell radius, can be written:

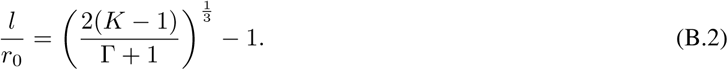

From equations (B.1) and (B.2), it is evident that, in the absence of functional grading, the strain inversion zone exists only if 6*κ*_*ECM*_ – 4*μ*_*ECM*_ < 9*κ*_1_. For *E*_*ECM*_ ≫ *E*_*cell*_ and *v*_*cell*_ < 1/2, strain inversion occurs for all *v*_*ECM*_ < 0.

## C Scaling of peri-cellular dilation

With *u*(*r*) solved in section A, the strain field was found from:

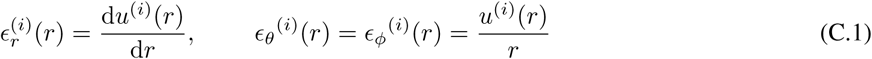

The dilatation (change in volume per unit volume), obtained from 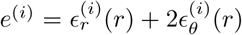, was constant within a layer with constant material properties. For a two-layer model consisting of the cell and ECM, the dilatation was calculated as:

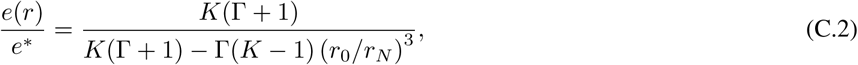

where *K* = *κ*_*ECM*_/*κ*_*cell*_ and Γ = (4/3)(*μ*_*ECM*_/*κ*_*cell*_), in which *κ*_*cell*_ and *κ*_*ECM*_ represent the bulk moduli of the cell and ECM, respectively, and *μ*_*ECM*_ is the shear modulus of the ECM. *e*^*^ = *p*/*κ*_2_ is the dilatation of the ECM in the absence of a cell. Clearly, *e*(*r*)/*e*(∞) = 1 for a spatially homogeneous ECM.

